# Mega-analysis of 31,396 individuals from 6 countries uncovers strong gene-environment interaction for human fertility

**DOI:** 10.1101/049163

**Authors:** Felix C. Tropf, Renske M. Verweij, Peter J. van der Most, Gert Stulp, Andrew Bakshi, Daniel A. Briley, Matthew Robinson, Anastasia Numan, Tõnu Esko, Andres Metspalu, Sarah E. Medland, Nicholas G. Martin, Harold Snieder, S. Hong Lee, Melinda C. Mills

**Affiliations:** Department of Sociology/ Nuffield College, University of Oxford, Oxford OX1 3UQ, UK; Department of Sociology/Interuniversity Center for Social Science Theory and Methodology, University of Groningen, Groningen 9712 TG, The Netherlands; Department of Epidemiology, University of Groningen, University Medical Center Groningen, Groningen 9700 RB, Netherlands.; Department of Population Health, London School of Hygiene and Tropical Medicine, London WC1E 7HT, UK; The University of Queensland, Queensland Brain Institute, Brisbane, QLD 4072, Australia; Department of Psychology, University of Illinois at Urbana-Champaign, Champaign 61820-9998, USA; Department of Medical Epidemiology and Biostatistics, Karolinska Institutet, PO Box 281, Stockholm SE-171 77, Sweden; Estonian Genome Center, University of Tartu, Tartu, Estonia, 51010, 140 Cambridge 02142, MA, USA; Quantitative Genetics Laboratory, QIMR Berghofer Medical Research Institute, Brisbane, QLD 4029, Australia; School of Environmental and Rural Science, The University of New England, Armidale NSW 2351, Australia

**Author notes:** Corresponding author: Felix C. Tropf, Manor Rd, Ox1 3UQ.

**Keywords:** Fertility, Age at first birth, Gene-environment interaction, Missing heritability, Natural selection, GREML

## Abstract

Family and twin studies suggest that up to 50% of individual differences in human fertility within a population might be heritable. However, it remains unclear whether the genes associated with fertility outcomes such as number of children ever born (NEB) or age at first birth (AFB) are the same across geographical and historical environments. By not taking this into account, previous genetic studies implicitly assumed that the genetic effects are constant across time and space. We conduct a mega-analysis applying whole genome methods on 31,396 unrelated men and women from six Western countries. Across all individuals and environments, common single-nucleotide polymorphisms (SNPs) explained only ~4% of the variance in NEB and AFB. We then extend these models to test whether genetic effects are shared across different environments or unique to them. For individuals belonging to the same population and demographic cohort (born before or after the 20^th^ century fertility decline), SNP-based heritability was almost five times higher at 22% for NEB and 19% for AFB. We also found no evidence suggesting that genetic effects on fertility are shared across time and space. Our findings imply that the environment strongly modifies genetic effects on the tempo and quantum of fertility, that currently ongoing natural selection is heterogeneous across environments, and that gene-environment interactions may partly account for missing heritability in fertility. Future research needs to combine efforts from genetic research and from the social sciences to better understand human fertility.

**Authors Summary:** Fertility behavior – such as age at first birth and number of children – varies strongly across historical time and geographical space. Yet, family and twin studies, which suggest that up to 50% of individual differences in fertility are heritable, implicitly assume that the genes important for fertility are the same across both time and space. Using molecular genetic data (SNPs) from over 30,000 unrelated individuals from six different countries, we show that different genes influence fertility in different time periods and different countries, and that the genetic effects consistently related to fertility are presumably small. The fact that genetic effects on fertility appear not to be universal could have tremendous implications for research in the area of reproductive medicine, social science and evolutionary biology alike.

## Introduction

Twin and family studies from Western countries show that genetic factors may explain up to 50% of the differences in human fertility outcomes such as number of children ever born (NEB) or age at first birth (AFB) within a population [1–8]. It remains unknown, however, whether the same genes are important for fertility across different environments or whether gene-environment interaction modifies genetic effects on fertility. This is a vital question for at least three reasons. First, the most successful and widely-used design to detect the approximate location of genetic variants associated with complex traits is a meta-analysis of genome-wide association studies (GWAS) from multiple populations [9]. This approach assumes genetic effects on a trait to be universal across environments. However, concerning fertility, this requires investigation given that environmental upheavals such as the introduction of the pill or educational expansion have substantially changed fertility behavior in the recent past [10,11]. A second and interrelated point is that studies resorting to molecular genetic data to quantify heritability as the variance in a trait explained by genetic variance result in lower estimates than family studies [12] – and this is true also in fertility research [1,2,7,13,14]. This discrepancy might, amongst other reasons, be a consequence of the interaction between environment and genes. Family studies are conducted amongst members of the same populations, whereas for example GWAS use data from individuals across populations. If genes can explain variance in fertility within but not between populations, heritability estimates based on different populations will be smaller than within populations [12,15]. Third, Fisher’s fundamental theorem of natural selection predicts at environmental equilibrium (close to) zero additive genetic effects on fitness-related traits such as fertility, because genes that reduce fitness are expected to have been passed on to the next generation to a lesser extent [16]. Nevertheless, additive genetic influences on fertility are well established and a potential explanation is that the genes that are important for fertility differ across environments [17].

Twin and family designs cannot be used to answer the question as to whether different genes are important for fertility across populations or birth cohorts since relatives usually live in the same country and twins always have the same age. However, with the advent of molecular genetic data and complementary analytical techniques and software, it has become possible to examine the genetic material of unrelated individuals across different (historical) populations and therefore the unique possibility exists to test whether the same genes influence a trait across diverse environments [18–22]. In this study, we exploit these advances for the first time, by empirically assessing whether genetic effects on fertility differ across geographical and historical environments.

We pooled a series of large datasets consisting of 31,396 unrelated (~ second cousin, IBS<0.05, see Material and Methods) genotyped men (n = 10,489) and women (n = 20,907) from six countries and seven study populations for analysis (for the US: HRS, ARIC; for the Netherlands: LifeLines; For Sweden: STR/SALT; for Australia: QIMR; for Estonia: EGCUT; for the UK: TwinsUK) who are assumed to have completed their reproductive period (*age*_*men*_ > 50; *age*_*women*_ > 45). We first conducted a mega-analysis, which is based on individual information from different populations in contrast to a meta-analysis that uses summary statistics of analyses conducted within populations, and applied whole genome methods [20,21] using GCTA software [18] to estimate SNP-heritability 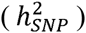. SNP-heritability is the proportion of total phenotypic variance that is explained by common genome-wide SNPs. Based on a previous study using data from women from the Netherlands and the UK, we expect 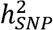 to be around 0.10 for number of children ever born and around 0.15 for AFB [23].

Second, to investigate gene-environment interaction, we follow two strategies: the first one consists in fitting multiple genetic relatedness matrices in our model, one global matrix for all individuals and more matrices indicating whether individuals lived in the same population and/or were part of the same birth cohort. The global matrix estimates the effects genes have across all environments, whereas the population/birth cohort specific matrices estimate context specific genetic effects [see 18 and Material and Methods for our specifications]. The second strategy consists in fitting bivariate genetic models to investigate the moderating effect of the postponement transition on genetic effects on fertility [see 22,24 and Material and Methods for our specifications]. This model allows us to estimate 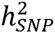 separately for different birth cohorts as well as the genetic correlation across them. To maximize power in these models, we divided all populations into two demographic birth cohorts. A central turning point in the reproductive environment of the 20^th^ century occurred when AFB experienced a massive postponement of up to 4-5 years in nearly all advanced societies, the so-called ‘postponement transition’ [25], or Second Demographic Transition [10,11,26,27]. The primary reasons proposed for fertility postponement have been women’s increased educational attainment and their employment in the labour force, triggered by factors such as the availability of effective contraception [10,11]. Cultural transformations in terms of sexual freedom, family planning and the timing and role of children are also central [26,27]. To investigate the moderating effect of fertility postponement we divide individuals into birth cohorts born either before or after this massive postponement in AFB in the past century [10,11,25,28].

## Results

### Descriptive findings

The descriptive statistics for NEB and AFB for all populations under study (LifeLines, TwinsUK, STR, Estonia, HRS, ARIC and QIMR) as well as the pooled data separate for men and women can be found in S1 Table. The participants were born between 1903 and 1967. The mean number of children per woman is 2.0 in Estonia, Sweden and the UK and 3.3 in Australia. For men, the lowest reported number of children is in Sweden and Estonia with around 1.9 children per man and is the highest in Australia at around 3.4. AFB was available for the Netherlands, UK, Sweden, Estonia and Australia. For both men and women, it was lowest in Estonia with an average of 24.6 for women and 27.7 for men and highest in Australia with 26.7 and 29.8 for men. Individuals who start reproducing at a later age have fewer children, with correlations between NEB and AFB ranging between −0.24 (Netherlands) and −0.38 (Australia; S2 Table). This pattern is less consistent across countries; for example in Australia, the highest fertility levels are observed, despite having the highest AFB. This reflects heterogeneity in fertility levels across countries with Australia having traditionally higher fertility levels than other Western countries (for a trend comparison of the total fertility rate across countries see S1 Fig).

### Demographic Trends

Fig 1 shows the trends in AFB during the 20^th^ century for the countries in our study based on population data if available (see Material and Methods for details). We observe the well-established U-shaped pattern of AFB of a falling AFB in the first half of the 20^th^ century followed by a turning point and upturn in the trend of AFB towards older ages. This postponement transition in fertility timing was accompanied by a strong drop in completed fertility in most countries [29].

**Figure 1.**
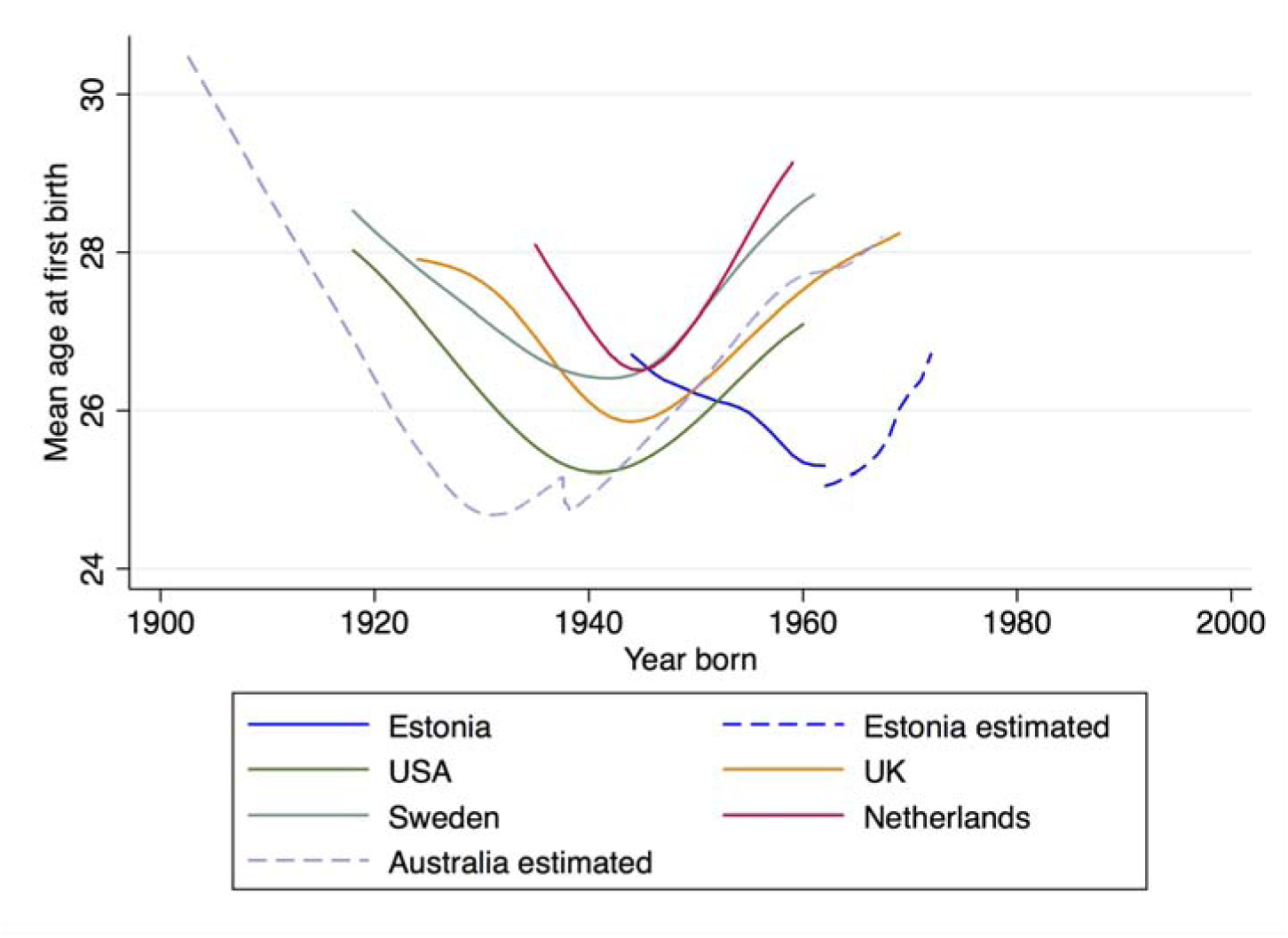
**Trends in age at first birth in cohorts from the US, UK, Sweden, the Netherlands, Estonia and Australia (1903-1970)**.

*Note*: Trends in the mean age at first birth are moving averages based on aggregated data obtained from Human Fertility Database and the Human Fertility Collection (for details see Material and Methods). For Australia, no official data has been available and the trends have been estimated from the QIMR dataset. See Supplementary Fig. S2 for the birth cohort specific average in the QIMR data.

Sociocultural and technological changes, such as the introduction of effective contraception, educational expansion or changing norms in reference to sexuality and family planning, have largely driven these trends [10,11]. These environmental changes occurred in specific time periods in each country. In order to test for gene-environment interaction in our analyses, we split the data into birth cohorts born before and after the turning point of fertility postponement to reduce environmental heterogeneity amongst the individuals who are members of the same birth cohort. This turning point differs across countries (Fig 1) with Australia having the earliest start of postponing (1939) and Estonia the latest (1962; see S3 Table for all turning points and details). Differences in the onset of the postponement transition are well established and can be due to, for example, political reasons. This is the case of Estonia, for example, where early AFB had been strongly promoted by political incentives when it still was part of the Soviet Union in time periods prior to 1990 [30].

### Genetic effects on fertility from the whole genome

#### Model 1: SNP heritability of AFB and NEB across environments

Not taking environmental differences into account, SNP based heritability (h^2^_SNP_) is significant and low for number of children ever born and age at first birth (Table 1). For NEB, h^2^_SNP_ is 0.038 (SE = 0.0097, p-value = 2.0x10^−5^) and for AFB it is 0.053 (SE = 0.019 p-value = 0.0020; these estimates are based on the full genetic relatedness matrix - see Material and Methods). These findings mean that around four per cent of the variance in NEB and around five per cent in AFB can be attributed to common, additive genetic effects in the pooled data. These estimates are much lower than those reported in other studies [23].

**Table 1.**
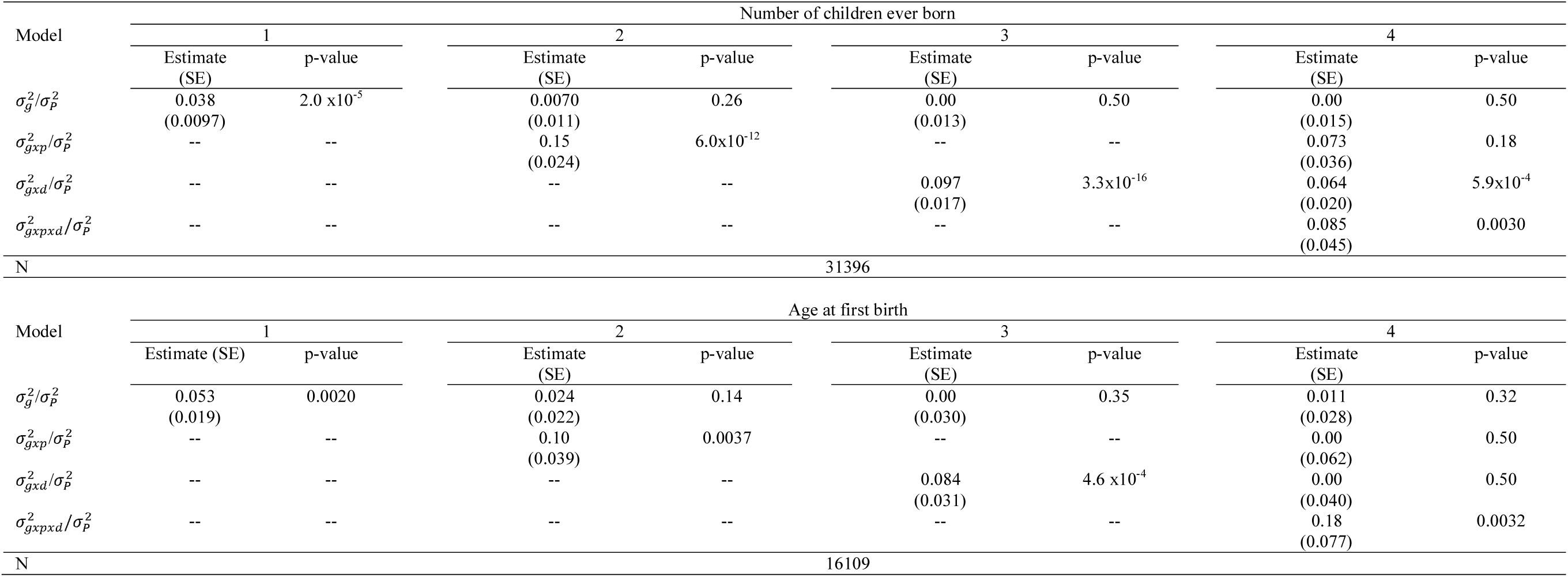
**Heritability estimates of the full GREML model and gene environment interaction models for number of children ever born (NEB) and age at first birth (AFB)**

| Model | Number of children ever born |  |  |  |  |  |  |  |
| --- | --- | --- | --- | --- | --- | --- | --- | --- |
|  | 1 |  | 2 |  | 3 |  | 4 |  |
|  | Estimate (SE) | p-value | Estimate (SE) | p-value | Estimate (SE) | p-value | Estimate (SE) | p-value |
| $\sigma_g^2/\sigma_P^2$ | 0.038<br>(0.0097) | $2.0 \times 10^{-5}$ | 0.0070<br>(0.011) | 0.26 | 0.00<br>(0.013) | 0.50 | 0.00<br>(0.015) | 0.50 |
| $\sigma_{gxp}^2/\sigma_P^2$ | -- | -- | 0.15<br>(0.024) | $6.0 \times 10^{-12}$ | -- | -- | 0.073<br>(0.036) | 0.18 |
| $\sigma_{gxd}^2/\sigma_P^2$ | -- | -- | -- | -- | 0.097<br>(0.017) | $3.3 \times 10^{-16}$ | 0.064<br>(0.020) | $5.9 \times 10^{-4}$ |
| $\sigma_{gxpxd}^2/\sigma_P^2$ | -- | -- | -- | -- | -- | -- | 0.085<br>(0.045) | 0.0030 |
| N | 31396 |  |  |  |  |  |  |  |
| Model | Age at first birth |  |  |  |  |  |  |  |
|  | 1 |  | 2 |  | 3 |  | 4 |  |
|  | Estimate (SE) | p-value | Estimate (SE) | p-value | Estimate (SE) | p-value | Estimate (SE) | p-value |
| $\sigma_g^2/\sigma_P^2$ | 0.053<br>(0.019) | 0.0020 | 0.024<br>(0.022) | 0.14 | 0.00<br>(0.030) | 0.35 | 0.011<br>(0.028) | 0.32 |
| $\sigma_{gxp}^2/\sigma_P^2$ | -- | -- | 0.10<br>(0.039) | 0.0037 | -- | -- | 0.00<br>(0.062) | 0.50 |
| $\sigma_{gxd}^2/\sigma_P^2$ | -- | -- | -- | -- | 0.084<br>(0.031) | $4.6 \times 10^{-4}$ | 0.00<br>(0.040) | 0.50 |
| $\sigma_{gxpxd}^2/\sigma_P^2$ | -- | -- | -- | -- | -- | -- | 0.18<br>(0.077) | 0.0032 |
| N | 16109 |  |  |  |  |  |  |  |

*Note*: 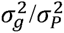 = proportion of observed variance in the outcome associated with genetic variance across all environments, 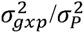 = proportion of observed variance in the outcomes associated with *additional* genetic variance within populations, 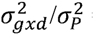 = proportion of observed variance associated with *additional* genetic variance within demographic birth cohorts, 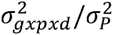 = proportion of observed variance associated with *additional* genetic variance within populations and demographic birth cohorts, p-values are based on likelihood-ratio test comparing the full model with the model with one constraining the particular effect to be zero, all analyses include the first 20 Principal Components, outcomes are standardized for sex, birth year and country.

#### Model 2: Genes x population interaction (g x p)

A potential reason to explain why estimates are lower than expected is that the SNPs important for fertility have different effects across environments. Model 2 therefore adds an interaction term to Model 1 that captures the influence of genetic variance on fertility only within populations. The gene-population interaction models for NEB and AFB show that shared genetic effects across populations are much lower than genetic effects within populations. With respect to NEB, the shared genetic effects across populations are negligible (0.0070, SE = 0.011, p-value = 0.26), whereas within populations additional additive effects are estimated to be 0.15 (SE = 0.024, p-value = 6.0 x10^−12^). The same applies to AFB, where shared genetic effects are estimated to be only 0.024 (SE = 0.022, p-value = 0.14), whereas the within population effect is 0.10 (SE = 0.039, p-value=0.0037; Table 1; Model 2). These results show that there is little overlap in SNPs that influence fertility across populations, and that most of the SNPs influencing fertility are population specific.

#### Model 3: Genes x demographic birth cohort (g x d)

Similar to the Model 2, in which we modeled population specific effects, we also examined whether there were genetic influences on fertility that were specific to birth cohorts. We find that there is additional genetic variance explanation for individuals who live in the same demographic cohort. While h^2^_SNP_ for all birth cohorts is estimated at zero for both NEB (SE = 0.013, p-value = 0.50) and AFB (SE = 0.03, p-value = 0.35), for individuals living in the same demographic cohort there is a significant additional genetic variance component of 0.097 (SE = 0.017, p-value = 3.3x10^−16^) for NEB and 0.084 (SE = 0.031, p-value = 4.6 x10^−4^) for AFB (Table 1; Model 3). Thus, similar to what we observed for the different populations, we find that SNPs influencing fertility traits are specific to cohort.

#### Model 4: Genes x population x demographic birth cohort (g x p x d)

Including a gene-environment interaction term that takes into account both the population and the demographic cohort simultaneously (Model 4), we observe that for NEB the interaction with demographic cohort (0.064, SE = 0.020, p-value = 5.9x10^−4^) and the interaction with population and demographic cohorts (0.085, SE = 0.045, p-value = 0.0030) are significant. This suggests that living in the same demographic cohort increases h^2^_SNP_ independent of whether individuals live in the same population, but rather living in the same population and the same demographic period additionally increases h^2^_SNP_. For AFB h^2^_SNP_ is only significantly different from zero for individuals living in the same population and demographic cohort (0.18, SE = 0.077, p-value = 0.0032).

### Overall SNP based heritability for each model

Subsequently, overall heritability estimates were calculated as the sum of the different components of each model to examine the increase in heritability estimates when including the different interaction terms (See Fig 2 corresponding to S4 Table). The overall h^2^_SNP_ for NEB increases almost fivefold, from 0.04 (SE = 0.01; Model 1) to 0.22 (SE = 0.026), when population and demographic cohort are taken into account. For AFB, the trend is very similar, with h^2^_SNP_ of 0.053 (SE = 0.019) in the baseline Model 1 and 0.19 (SE = 0.039) in the genes x population x demographic cohort interaction model, when population and demographic cohort are taken into account.

**Figure 2.**
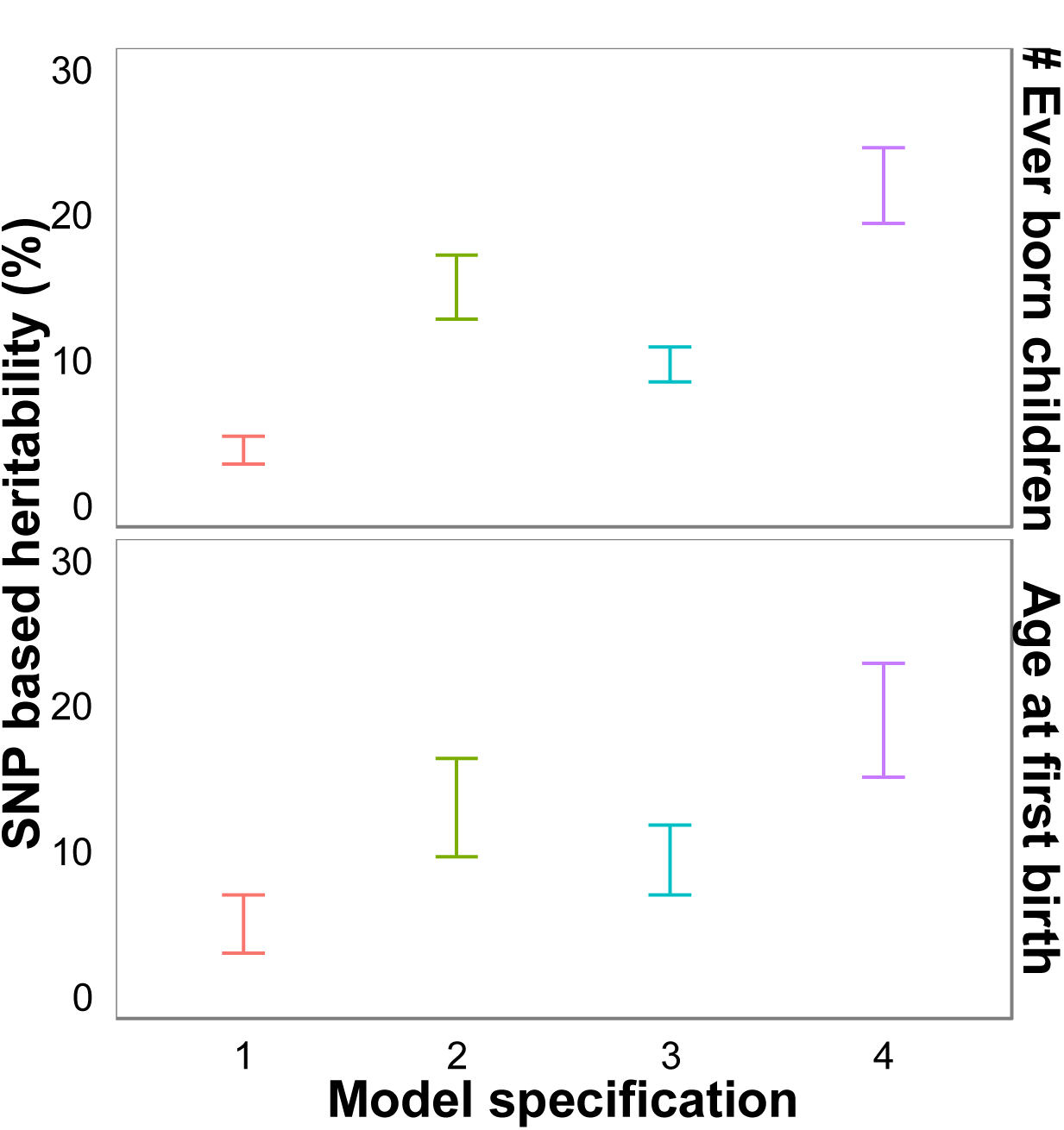
**Bar Charts of the SNP-heritability estimates in number of children ever born (NEB) and age at first birth (AFB) for the different model specifications from Table 1**.

*Note*: SNP-heritability as the sum of genetic variance over the total variance in Model specification 1 = amongst all individuals, 2 = amongst individuals living within the same population, 3 = amongst individuals living within the same demographic birth cohort born either before or after fertility postponement, 4 = amongst individuals living in the same population and demographic birth cohort, dots = estimate, lines = estimate ± 1 SE, The corresponding table to Figure 2 an be found in Supporting Table S4.

### Sensitivity analysis: Genes x Sex

The analyses presented are based on pooled datasets of men and women. However, two data sources contain (almost) only women (TwinsUK and ARIC). To the extent that different genes influence fertility across sexes, this might drive the observed differences across populations. We therefore conducted a sensitivity analysis extending Model 3 to a genes x population x sex interaction model. We find that considering sex-differences does not significantly improve the model fit (p-value for AFB 0.5, for NEB 0.093) and therefore are confident that our findings do not result from sex-differences (S6 Table).

### Bivariate analysis

We complementarily estimated a bivariate model based on Model 2 and splitting data for demographic cohort (see Material and Methods), which allows us to estimate genetic effects across 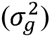 and within 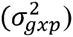 populations separately for different demographic birth cohorts and investigate whether genetic effects are correlated across demographic birth cohorts. Table 2 shows that 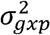 estimates for NEB within populations are significant for both demographic cohorts before (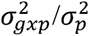, SE = 0.039, p-value = 9.6x10^−6^) and after (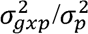, SE = 0.048, p-value = 0.0010) fertility postponement. It furthermore shows a positive correlation of genetic effects on NEB across demographic cohorts within populations (1.00, SE = 0.35, p-value = 1.3x10^−5^). In Model 4 of Table 1, this remained suggestive, since the genetic effects within populations but shared across demographic cohorts 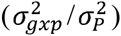 were non-significant (0.073, SE = 0.036, p-value = 0.18). The bivariate model for the AFB finds some evidence that in both demographic cohorts genetic effects are observed (before fertility postponement 0.099, SE = 0.073, p-value = 0.083; after fertility postponement 0.13, SE = 0.074, p-value = 0.070), although these effects were marginally significant. However, there is no evidence that genetic effects correlate across demographic birth cohorts (0.11, SE = 0.59, p-value = 0.27), which is well in line with the null-estimate 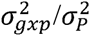 (0.00, SE = 0.062, p-value = 0.50).

**Table 2.**
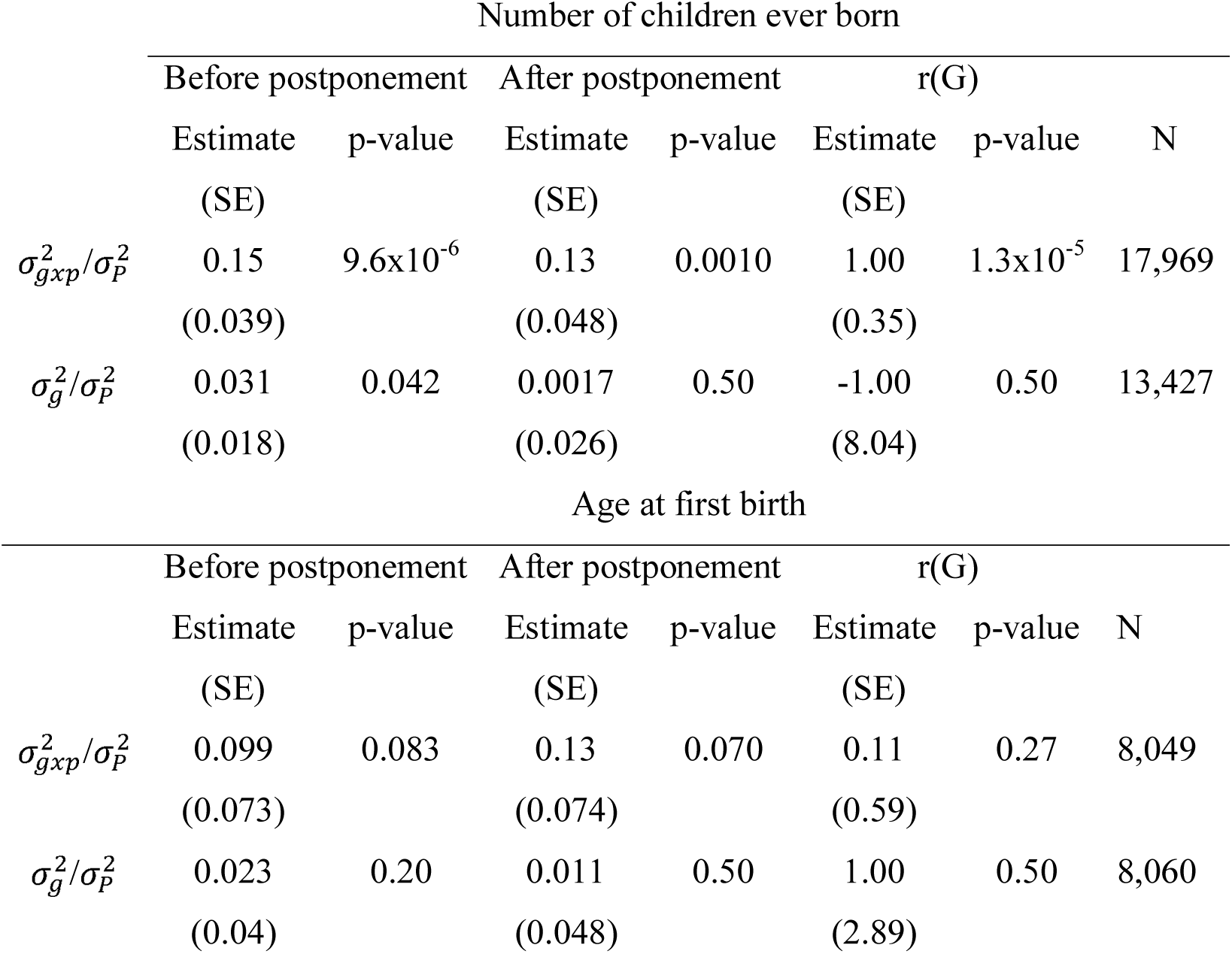
**Bivariate analysis of Model 2 to estimate genetic correlations for gxp (genes x population) or global g (gene) component before and after fertility postponement**

*Note*: 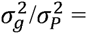 proportion of observed variance in the outcome associated with genetic variance across all environments, 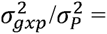 proportion of observed variance in the outcomes associated with *additional* genetic variance within populations, 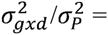 proportion of observed variance associated with *additional* genetic variance within demographic birth cohorts, r(G) = genetic correlation, p-values are based on likelihood-ratio test comparing the full model with the model with one constraining the particular effect to be zero, all analyses include the first 20 Principal Components, outcomes are standardized for sex, birth year and country.

Genetic effects shared across all populations 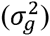 are only significant for NEB and birth cohorts born before fertility postponement (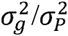, SE = 0.018, p-value 0.042) while non-significant for younger cohorts and for AFB. Genetic correlations are therefore not interpreted.

## Discussion

Using data from seven populations and six countries, we demonstrate that genetic effects on fertility outcomes–number of children ever born (NEB) and age at first birth (AFB) – differ across temporal and spatial environments. For NEB, genetic effects within populations are stronger than across populations, but correlate between individuals who were born before or after the turning point in fertility postponement of the 20^th^ century. For AFB, genetic effects are only significant if individuals live in the same demographic cohort and the same population. The full gene-environment interaction model (Model 4) as well as the bivariate analyses provide no evidence for shared genetic effects for each phenotype across populations and demographic birth cohorts. Our results show that different SNPs are associated with fertility traits in different populations and birth cohorts, and there are hardly any genetic effects that are consistently related to these traits across populations and cohorts. Our results uncover a strong interplay of genetic and environmental factors influencing human fertility.

Quantitative geneticists have been puzzled by low heritability estimates based on GWAS findings or even whole-genome estimates such as GREML model as we apply it in the current study, describing the phenomenon of ‘missing heritability’ [12]. Previous attempts to explain missing heritability partly by non-additive genetic effects remain empirically untested [31] or find only little support [32]. Our findings of strong gene-environment interaction imply first that the detection of genetic variants associated with fertility traits is a major challenge using meta-analyses of GWAS on individuals from different populations. Likewise, predictions out of the discovery sample might be difficult, because discovered SNPs might have different effects in different samples. Second, they imply that lower heritability estimates from GWAS studies compared to GREML approaches or family studies might be due to the fact that genetic effects are (to some extent) not universal but context specific. In the model considering gene-environment interaction across population and demographic cohort, we report heritability findings of 0.22 for NEB and 0.19 for AFB (see Fig 2 and S4 Table), which are fourfold higher than across all contexts and approach heritability estimates from family models [2,14]. It is therefore central to understand the cultural and environmental factors that interact with human fertility as well as their origins across (family) environments in order, for example, to define missing heritability or validate the findings from twin studies. It is to be noted that our findings are probably fostered by the strong behavioural and social nature of fertility, which might be more sensitive to cultural and societal heterogeneity than for example morphological traits. A recent investigation by Yang et al. [33] shows that missing heritability for the anthropometric traits height and body mass index is negligible when using whole genomic sequencing data in a new GREML model and assuming that family models overestimate heritability.

Demonstrating that genetic effects on fertility outcomes differ across environments, our study substantially contributes to the current knowledge on the genetic architecture of human reproduction. Previous twin studies show for several countries and birth cohorts that fertility outcomes such as NEB and AFB are genetically influenced [1,2]. However, it remained unclear whether the same genes are associated with fertility across environments. Using molecular genetic data and GREML methods [18,20,21], we were able to relate the genetic material of individuals across environments and found that common SNPs explain substantially more variance within than between countries and birth cohorts for fertility traits.

Previous twin and family studies furthermore suggest that the level of heritability of fertility traits can change across time and space [5,7,34,35]. However, these differences could not be statistically validated. In the current study, we proposed a multi-matrix approach to test for gene-environment interaction but also applied bivariate GREML models across birth cohorts [22,24]. Bivariate GREML models allow estimating SNP-heritability within two independent samples was well as the genetic correlation across them. We cannot confirm the suggestion that the level of heritability changed over time, but find that heritability levels are comparable before and after the strong fertility postponement in the past century.

Different levels of heritability have also been reported across countries [2]. Our multi-matrix GREML approach distinguishes between pairs of individuals who are living in the same or in different populations. The resulting within population estimate is therefore an average across all populations and we cannot compare different levels of heritability across populations. A more desirable study design would be a multivariate genetic modelling approach as we presented it in a bivariate design to investigate differences across demographic birth cohorts. However, this approach was not possible in our study due to small sample sizes in each population and a consequent lack of statistical power [22], but might become feasible in the future with better data availability.

Our findings are of interest to scientists within the medical, biological and social sciences alike [1,2,36,37]. Research has successfully identified genetic variants associated with reproductive diseases and traits [37]. However, it remains unknown how these affect realized fertility. We find no evidence that genetic effects underlying fertility in one country predict fertility outcomes in another one. Genetic effects on fertility outcomes are rather strongly dependent on an individual’s environment. Recently, social scientists have made large efforts to integrate molecular genetics into their research [1,2,23,34,38–42]. However, when it comes to reproductive health, environmental factors are also likely to be critical in understanding how genetic factors are modified in relation to fecundity and infertility.

For evolutionary biologists, our findings have at least two important implications. First, the number of children ever born has been used as a proxy for fitness, given the diminishing child mortality rate in contemporary societies [4,23,36]. Additive genetic variance therefore indicates currently ongoing natural selection under environmental equilibrium within populations: if all else equal, genes that lead to a higher number of offspring will have a higher frequency in future generations. Due to natural selection, Fisher predicted additive genetic variance in fertility to be (close to) zero in the absence of gene environment interaction, since genes that reduce fitness are passed on to the next generation to a lesser extent thereby reducing their frequencies [16]. Nevertheless, we find significant additive genetic influences on fitness traits such as NEB and AFB – substantial yet lower than heritabilities observed for morphological traits such as height [14,15,23,43]. Finding significant genetic influences on this these proxies of fitness suggests that, along with sociocultural changes surrounding fertility, genetic variants under selection have also changed [for review see 1,for review see 2,5,7,17,34,for comment see 44–46]. Gene-environment interaction can explain why we find additive genetic variance in fitness related traits despite natural selection.

Second, previous research has uncovered an ongoing natural selection in contemporary societies [3,4,23,47,48] and even attempted to forecast changes in for example height and blood pressure across generations [4]. For valid evolutionary predictions about observable changes in traits across generations due to currently ongoing natural selection, fertility needs to be consistently heritable, the same genes need to be under selection across generations and the direction of the selection needs be similar. Our results demonstrate moderate genetic influences on fertility within populations indicating potentially ongoing human evolution. However, this potential is delimited in at least two ways: First, genetic effects on fertility strongly differ across countries and therefore may lead to heterogeneity across human populations rather than to universal changes in humans. Second, the finding that genetic effects underlying proxies of fitness vary so markedly across time periods suggests that substantial caution is needed when inferring long-term evolutionary predictions.

For social scientists, genetic influences had been originally thought of as biological constraints on human reproductive behavior [42]. Yet some previous studies showed that genetic predispositions may underlie decision making processes on fertility timing and motivation [6,7,49,50]. It has been suspected that genetically based behavioural and psychological traits have become more important than physiological ones in the recent past [6,8,34,51]. This hypothesis remains to be tested, but our results confirm that genetic influences on fertility have evolved with social changes in the reproductive environment and therefore underscore the necessity to integrate social factors into genetic research on fertility.

Overall, our study uncovers great challenges for investigations into the genetic architecture of fertility, which can only be overcome by interdisciplinary work between both social scientists and geneticists using ever larger datasets, with combined information from genetics and social sciences [36].

## Material & Methods

### Cohorts

In this study we combined data from seven cohorts and six countries. For the US, we use data from ARIC, HRS, for Estonia from EGCUT, for Australia QIMR data from the Australia Twin and Family Register, for the Netherlands the LifeLines Cohort Study, for the United Kingdom TwinsUK and for Sweden the STR. All studies have received ethical approval.

### ARIC

ARIC (Atherosclerosis Risk in Communities Study) is a community-based prospective cohort study of 15,792 adults, ages 45–64. Participants were identified by probability sampling from four U.S. communities (Forsyth County, North Carolina; Jackson, Mississippi; suburban Minneapolis, Minnesota; and Washington County, Maryland) and were enrolled between 1987 and 1989 [52–54].

### HRS

The Health and Retirement Study (HRS) is an ongoing cohort study of Americans, with interview data collected biennially on demographics, health behavior, health status, employment, income and wealth, and insurance status. The first cohort was interviewed in 1992 and subsequently every two years, with 5 additional cohorts added between 1994 and 2010. The full details of the study are described in [55].

### EGCUT

Estonian data come from of the Estonian Genome Center Biobank, University of Tartu (EGCUT, www.biobank.ee), a population-based database which comprises the health, genealogical and genome data of currently more than 51,530 individuals [56]. Each participant filled out a Computer Assisted Personal Interview including personal data (place of birth, place(s) of living, nationality etc.), genealogical data (family history, three generations), educational and occupational history and lifestyle data (physical activity, dietary habits, smoking, alcohol consumption, and quality of life).

### QIMR

Data for Australia was received from the Queensland Institute for Medical Research (QIMR). The participants were drawn from cohorts of adult twin families that have taken part in a wide range of studies of health and well-being via questionnaire and telephone interview studies, and recruitment was extended to their relatives (parents, siblings, adult children and spouses).

### LifeLines Cohort Study

The LifeLines Cohort Study [57] is a multi-disciplinary prospective population-based cohort study from the Netherlands, examining in a unique three-generation design the health and health-related behaviours of 167,729 persons living in the North of The Netherlands including genotype information from more than 13,000 unrelated individuals. It employs a broad range of investigative procedures in assessing the biomedical, socio-demographic, behavioural, physical and psychological factors which contribute to the health and disease of the general population, with a special focus on multi-morbidity and complex genetics.

### TwinsUK

For the UK, we use data from TwinsUK, the largest adult twin registry in the country with more than 12,000 respondents [58]. The TwinsUK Study recruited white monozygotic (MZ) and dizygotic (DZ) twin pairs from the TwinsUK adult twin registry, a group designed to study the heritability and genetics of age-related diseases (www.twinsuk.ac.uk). These twins were recruited from the general population through national media campaigns in the UK and shown to be comparable to age-matched population singletons in terms of clinical phenotype and lifestyle characteristics.

### STR

The Swedish Twin Registry (STR) was first established in the late 1950s to study the importance of smoking and alcohol consumption on cancer and cardiovascular diseases whilst controlling for genetic propensity to disease. Between 1998 and 2002, the STR conducted telephone interview screening of all twins born in 1958 or earlier regardless of gender composition or vital status of the pair. This effort is known as Screening Across the Lifespan Twin study (SALT). A subsample of SALT (≈10,000) was genotyped as part of the TwinGene project [59] and we use the this information in the current study.

### Fertility trends

Aggregate data to describe country specific fertility trends have been obtained from the Human Fertility Database (HFD, http://www.humanfertility.org/cgi-bin/main.php) and the Human Fertility Collection (HFC, http://www.fertilitydata.org/cgi-bin/index.php) if available. Both data collections are joint projects of the Max Planck Institute for Demographic Research (MPIDR) in Rostock, Germany and the Vienna Institute of Demography (VID) in Vienna, Austria. The projects provide access to detailed and high-quality data on period and cohort fertility. The HFD is entirely based on official vital statistics. The HFC incorporates a variety of valuable fertility data from diverse, not necessarily official, data sources. All data are freely available after registration. We focused on fertility information for cohorts that was aggregated for individuals older than 45.

For the UK, official data on birth order have only been collected within marriage, and might be biased. We therefore relied on estimates from the Office for National Statistics, Cohort fertility, Table 2. Available at: http://www.ons.gov.uk/ons/publications/re-reference-tables.html?edition=tcm%3A77-2631333. For Estonia, data on completed reproduction by age 45 was only available until the year 1962. For subsequent cohorts, however, there was an estimate of AFB available based on official statistics at the age of 40. For Australia, no official data on a time series of cohort specific AFB was available and the trends are based on the data used for analysis in this study.

### Genotypes

We received genotype data from all cohorts, which we imputed according to the 1000 genome panel – except for Twins UK from which we already received the imputed data.

### Genetic-relatedness-matrix (GRM)

To estimate the genetic relatedness-matrix (GRM) the HapMap3 imputation panel was used as a reference set as it was optimized to capture common genetic variation in the human genome [60].We selected HapMap3 SNPs with an imputation score larger than 0.6. For quality control (QC), we excluded the SNPs with a larger missing rate than 5% after merging, lower minor allele frequency than 1% and which failed the Hardy-Weinberg equilibrium test for a threshold of 10^−6^. We merged the datasets subsequently applying QC again after merging each data set. 847,278 SNPs could be utilized to estimate the GRM between individuals. We used the software Plink [19] for the quality control and merging of the datasets and GCTA [18] to estimate the genetic relatedness matrix

The GRM *A*_*ik*_ is estimated for each pair of individuals j and k:

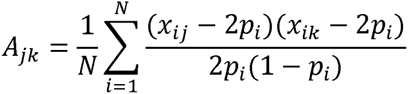

where *x*_*ij*_ and *x*_*ik*_ is the number of copies of the reference allele for the *i*^th^ SNP of the *j*^th^ or *k*^th^ individual and *p*_*i*_ is the frequency of the reference allele and *N* the number of SNPs. If two individuals had a higher genetic relatedness than 0.05, one was excluded from the analyses to avoid bias due to environmental confounders amongst close relatives.

### Phenotypes

Number of children ever born was available for all cohorts, but in ARIC and TwinsUK, however, only for women. NEB measures number of children a woman has given birth to or a man has fathered. It was either directly asked or we constructed it from questions on the date of birth of each child.

The measure is not perfectly harmonized across cohorts because some questionnaires explicitly exclude still-births (HRS, ARIC) while others remain undefined (TwinsUK asked in some waves: “How many children have you given birth to?”; EGCUT asked: “Do you have any biological children?”, and subsequently: “Fill in their names, gender and date of birth). In STR, Life Lines, QIMR as well as most of the waves of the TwinsUK, information on both the date of birth and death of the child was asked. In LifeLines and TwinsUK, we compared the live birth measure with number of children ever born and, as expected, given the diminishing mortality rate in both datasets, less than 0.2% of the children had not reached reproductive age and the correlation of number of children ever born and number of children reaching reproductive age was >0.98. We therefore expect no large bias due to the fact that in some countries still-births are excluded.

The questionnaires were furthermore heterogeneous in the maximum number of children that could be named. However, within each cohort, the maximum number of children has never been named more often than in 0.5 per cent of the cases and we do not anticipate that our results are influenced by this.

Information on AFB was available in all cohorts except for ARIC and the HRS. It was asked directly ( in Twins UK) or was constructed using the date of birth of the oldest child and the year of birth of the respondent.

Since fertility is strongly age dependent, we focus on women only with completed fertility history in reference to the phenotype. In general, the end of women’s reproductive lifespan occurs around the age of 45 and for men at the age of 50, thus, we only included individuals beyond those ages in our analyses. Furthermore, in vitro fertilization (IVF) – often related to twinning and multiple births – can bias results if IVF compensates genetically based infertility. However, in our TwinsUK sample, only 60 women reported using IVF, who we did not include in the final analyses.

For all models, both phenotypes were standardized (Z-transformed) by cohort, year of birth and sex.

## GREML Models

### GREML Models Common SNP heritability estimates (Model 1)

The genetic component underlying a trait is commonly quantified in terms of SNP-heritability 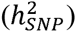 as the proportion of the additive genetic variance explained by common SNPs across the genome over the overall phenotypic variance 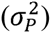 of the trait:

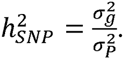

The phenotypic variance is the sum of additive genetic and environmental variance, i.e. 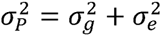 where 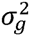 is the additive genetic variance explained by all SNPs across the genome and 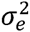 is the residual variance.

The methods we applied have been detailed elsewhere [18,20–22,24]. Briefly, we applied a linear mixed model

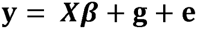

where **y** is an Nxl vector of dependent variables, N is the sample size, **β** is a vector for fixed effects of the overall mean (intercept), **g** is the Nxl vector with each of its elements being the total genetic effect of all SNPs for an individual, and **e** is an Nxl vector of residuals. The variance covariance matrix of the observed phenotypes is:

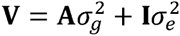

We have 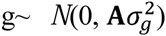 and 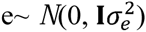, **A** is the genetic relationship matrix (GRM) estimated from SNPs and **I** is an identity matrix. The variance components are estimated using the restricted maximum likelihood (REML) approach.

### Genes x Population (Model 2)

The genes x demographic birth cohort interaction model is a joint model estimating global genetic effects for the fertility traits, effective between and within samples 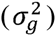 and the averaged additional genetic effects within cohorts 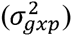:

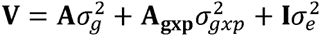

where **A** is the genetic relatedness matrix and **A**_**gxp**_ is a matrix only with values for pairs of individuals within populations 1-7:

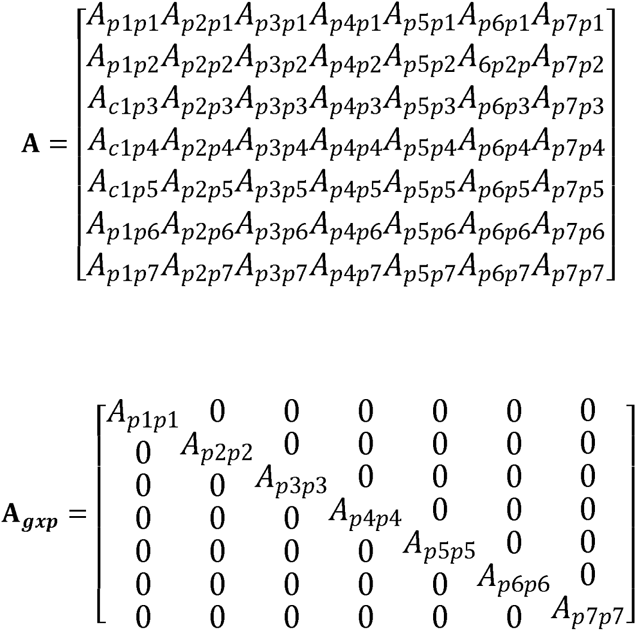

### Genes x Demographic birth cohort (Model 3)

The genes x demographic birth cohort interaction model is a joint model estimating the universal genetic effects for the traits, effective between and within samples 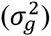 and the averaged additional genetic effects within defined birth cohorts 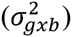:

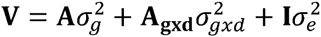

whereas **A** is the genetic relatedness matrix and **A**_**gxd**_ is a matrix only with values for pairs of individuals within the same demographic birth cohorts bl-2:

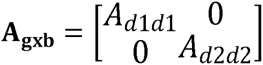

### Genes x Population x Demographic birth cohorts (Model 4)

Finally, we applied a model including both interaction terms from above and an additional interaction term **A**_**gxcxb**_ which is 0 for all pairs of individuals living in different time periods or in different cohorts

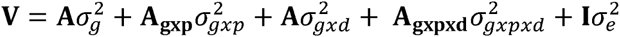

whereas **A** is the genetic relatedness matrix, **A**_**gxp**_ is a matrix only with values for pairs of individuals within populations 1-7 (Model 2), **A**_**gxd**_ is a matrix only with values for pairs of individuals within the same demographic periods b1-2 (Model 3) and **A**_**gxpxd**_ is a matrix only with values for pairs of individuals with both the same demographic periods and the same populations.

### Bivariate Model

For bivariate analyses [22,24], we split the data into individuals born before and after the turning point in fertility postponement in AFB (see also S5 Table). Based on Model 2, we estimate a bivariate model with two GRMs:

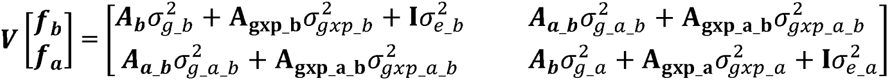

where as ***ƒ***_***b***_ **and *ƒ***_***a***_ are vectors of length *N*_*b*_ and *N*_*a*_ of fertility phenotypes (NEB or AFB) of individuals born **b**efore or **a**fter the postponement transition started, with N being the respective sample size of the subsets. Variance components refer to those from Model 2, whereas the lower index _b indicates that they are estimated in the subset of individuals born before and index a born after the start of the postponement transition. The index _b_a denominates the covariance of variances components across subsets. The correlation of the genetic factors are estimated as:

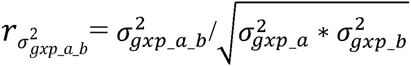

and

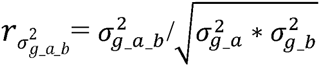

## Acknowledgement

Funding was provided by grants awarded to M.C.Mills: ERC Consolidator Grant SOCIOGENOME (615603), NWO (VIDI grant 452-10-012), ESRC/NCRM SOCGEN grant), ESRC/NCRM. The authors wish to thank the participants from all cohorts under study. We also wish to thank Patrick Präg, Mariana Bonnouvrier, Robert Hellpapp and Nicola Barban for useful comments on previous versions of this manuscript. For additional acknowledgments (participating cohorts), please see S1 text.

## Supporting Information (on request)

**Table S1.**
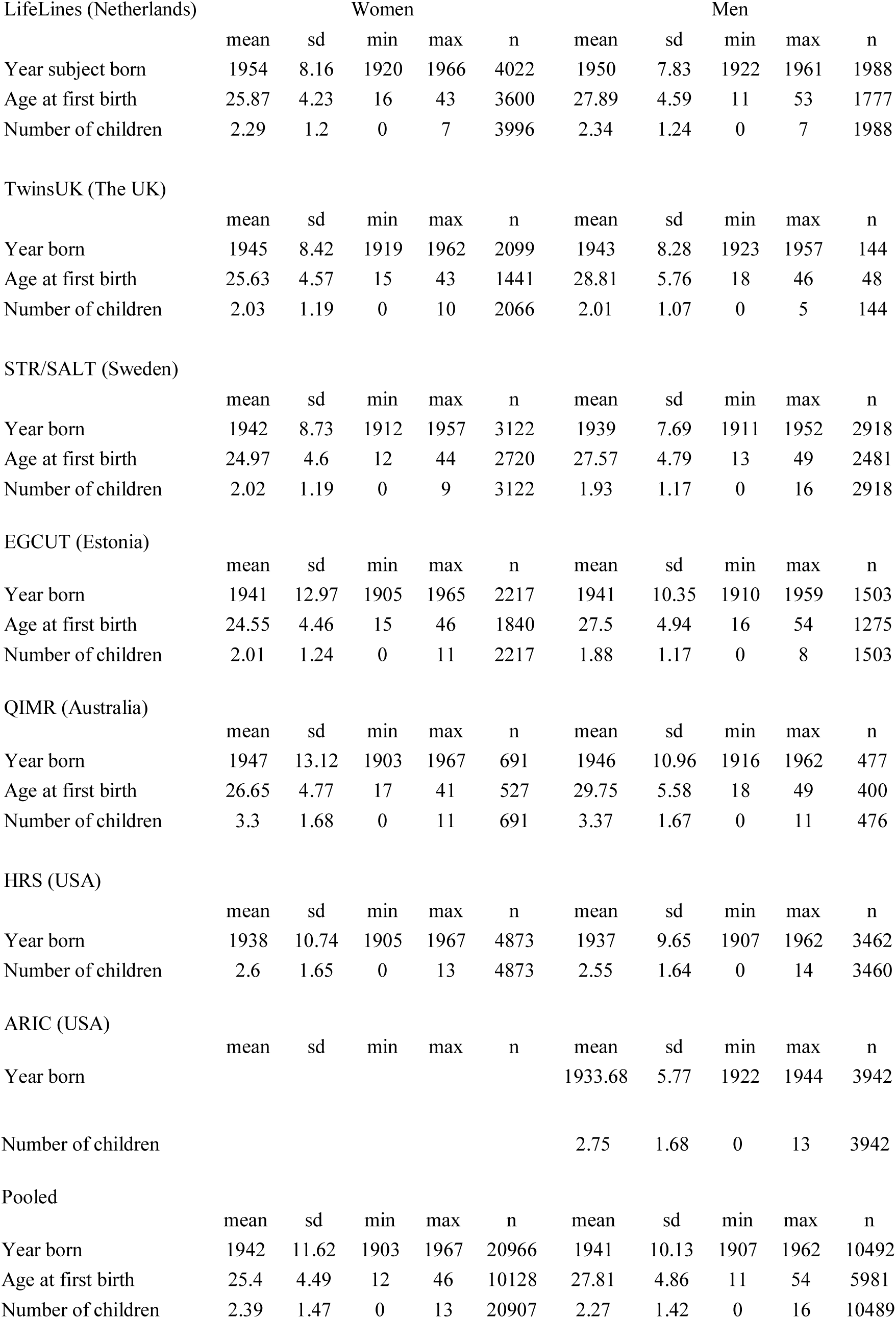
**Summary statistics for women and men for all datasets separately and pooled**

Note: The dataset descriptions can be found in the main text under Material & Methods.

**Table S2.** **Pearson’s correlation between AFB and NEB for each dataset containing both phenotypes**

|  | Correlation | Number of individuals |
| --- | --- | --- |
| LifeLines | -0.24 | 5351 |
| TwinsUK | -0.28 | 1456 |
| STR | -0.30 | 5201 |
| QIMR | -0.38 | 926 |
| EGCUT | -0.26 | 3115 |

**Fig. S1.**
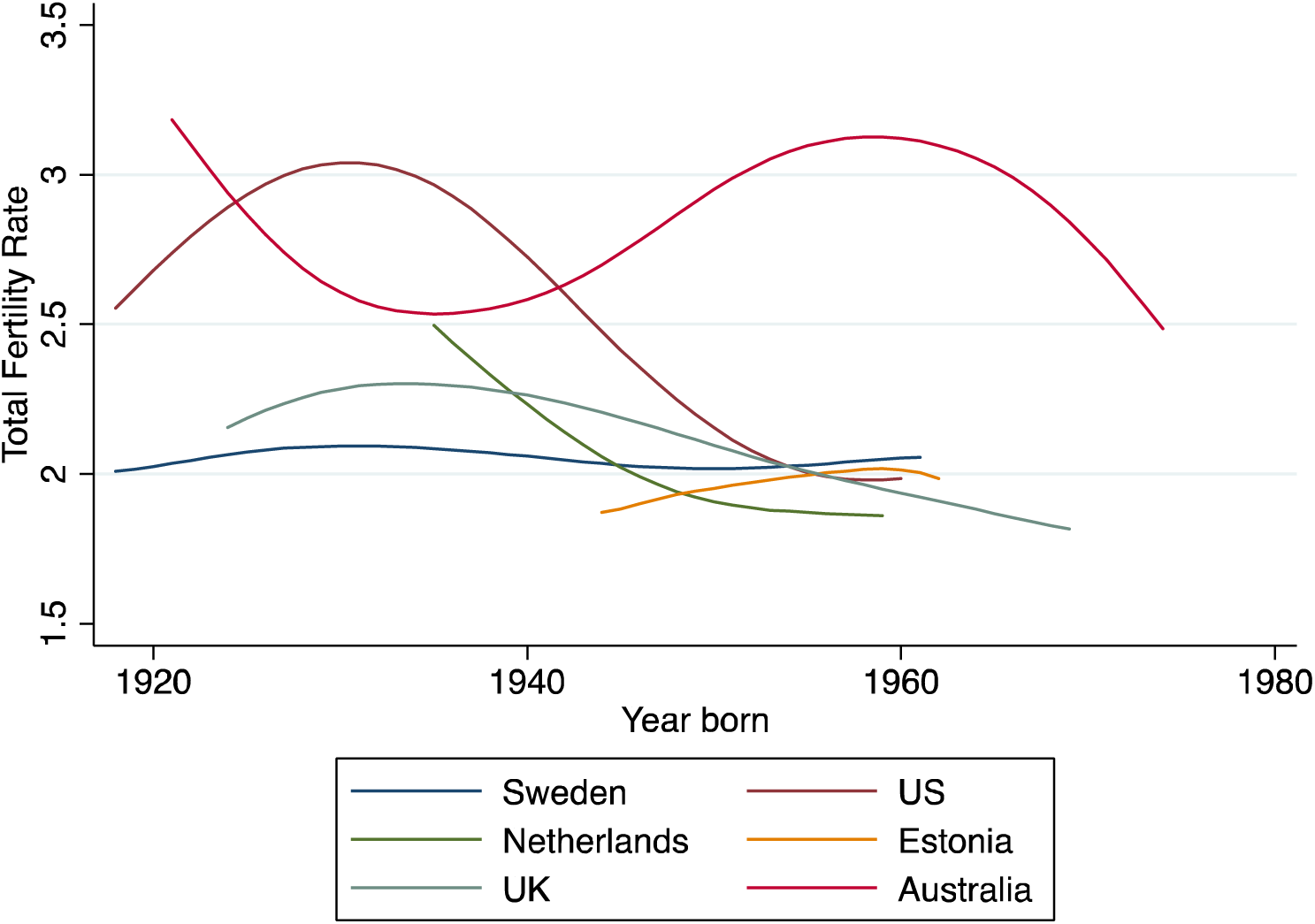
**Comparison of total fertility rates across countries under study**

Note: Aggregated data have been obtained from the Human Fertility Database and the Human Fertility Collection (for Australia; for details see Material and Methods in the main text). The Total Fertility rate refers to the expected average number of children born to a woman.

**Fig. S2.**
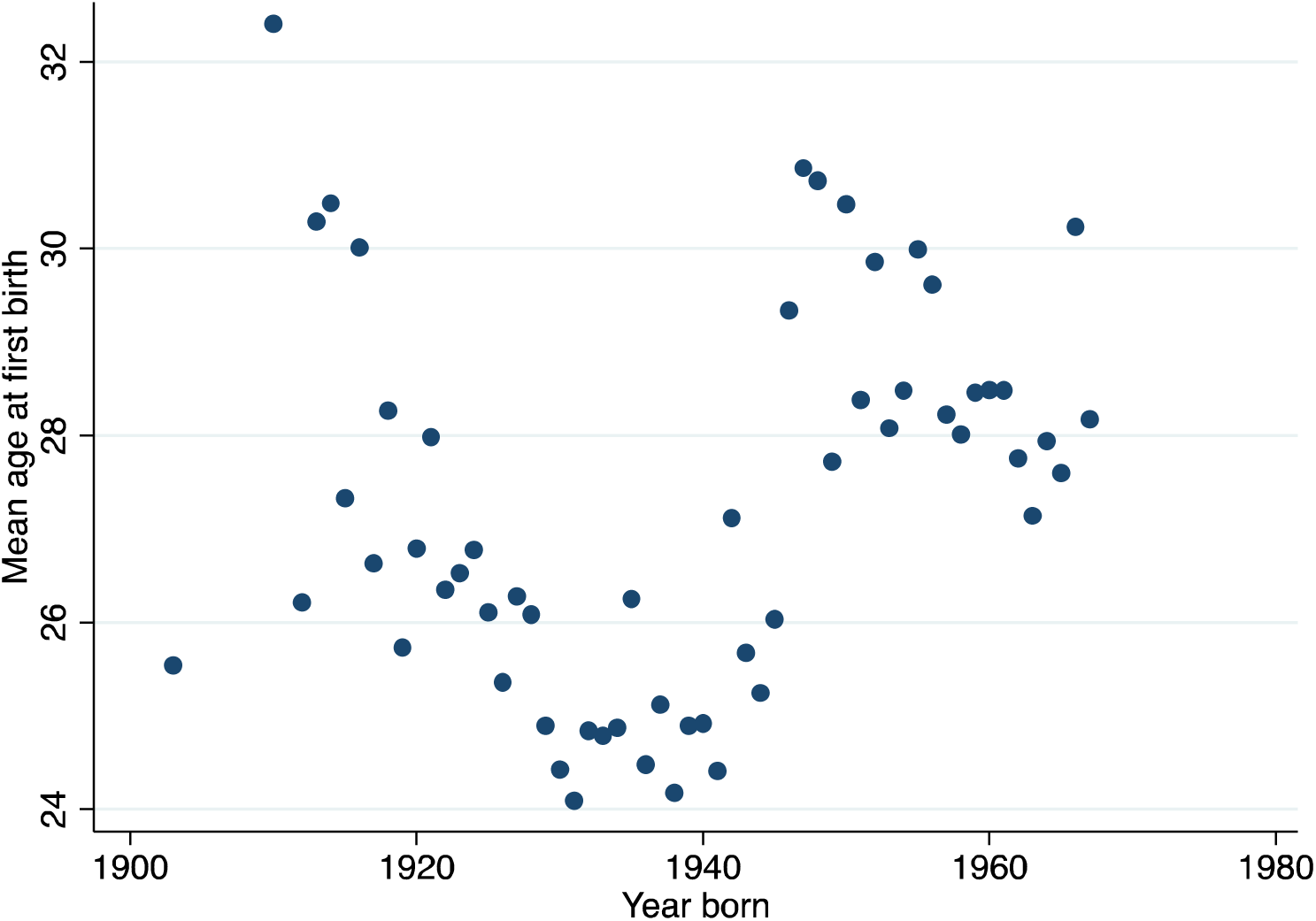
**Observed mean age at first birth in QIMR data for Australia**

Note: For a description of the data source QIMR see Material & Methods in the main text.

**Table S3.** **Observed low point in the age at first birth on the population level by country – indicating the turning point in the fertility postponement trend**

| Country | Turning point | Year of turning point | Data source <sup>a</sup> |
| --- | --- | --- | --- |
| Netherlands | 26.39 | 1944 | HFD |
| United Kingdom | 25.75 | 1944 | Official Statistics |
| Sweden | 26.39 | 1943 | HFD |
| Australia | 24.78 | 1939 | QIMR |
| Estonia | 25.05 | 1962 | HFD/HFC |
| United States | 24.63 | 1940 | HFD |

Notes: <sup>a</sup> See Material and Methods for details on data sources

Notes: ^a^ See Material and Methods for details on data sources

**Table S4.** **Overall SNP heritability estimated for Model specifications 1-4 for number of children ever born (NEB) and the age at first birth (AFB) as depicted in the main text Fig. 2**.

| Model | NEB |  |  |  | AFB |  |  |  |
| --- | --- | --- | --- | --- | --- | --- | --- | --- |
|  | 1 | 2 | 3 | 4 | 1 | 2 | 3 | 4 |
| SUM ($\sigma^2$)/$\sigma_p^2$ | 0.038 | 0.15 | 0.097 | 0.22 | 0.05 | 0.13 | 0.094 | 0.19 |
| SE | (0.0097) | (0.022) | (0.012) | (0.026) | (0.02) | (0.034) | (0.024) | (0.039) |

Note: SUM ( $\sigma^2$ )/ $\sigma_p^2$ refers to the total variance explanations from model specifications 1-4 (see Material and Methods)

Note: SUM 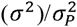 refers to the total variance explanations from model specifications 1-4 (see Material and Methods)

**Table S5.** **Sample sizes of datasets divided by demographic birth cohorts born before and after the onset of fertility postponement**

| Cohort | Before postponement | After postponement | Number of individuals |
| --- | --- | --- | --- |
| LifeLines | 1,039 | 4,971 | 6,010 |
| TwinsUK | 992 | 1,251 | 2,243 |
| ARIC | 3,373 | 569 | 3,942 |
| HRS | 4,972 | 3,361 | 8,333 |
| STR | 3,561 | 2,479 | 6,040 |
| QIMR | 343 | 825 | 1,168 |
| EGCUT | 3,699 | 21 | 3,720 |

Notes: See Material and Methods for details on data sources

Notes: See Material and Methods for details on data sources

**Table S6.** **Heritability estimates of the gene environment interaction models for population and sex by number of children ever born (NEB) and age at first birth (AFB)**

| Model | g x population x sex |  |  |  |
| --- | --- | --- | --- | --- |
|  | NEB |  | AFB |  |
|  | Estimate<br>(SE) | p-value | Estimate<br>(SE) | p-value |
| $\sigma_g^2/\sigma_P^2$ | 0.000001<br>(0.015) | 0.5 | 0.012<br>(0.032) | 0.35 |
| $\sigma_{gxp}^2/\sigma_P^2$ | 0.11<br>(0.036) | 0.0010 | 0.12<br>(0.059) | 0.013 |
| $\sigma_{gxs}^2/\sigma_P^2$ | 0.019<br>(0.020) | 0.18 | 0.021<br>(0.043) | 0.31 |
| $\sigma_{gxpxs}^2/\sigma_P^2$ | 0.051<br>(0.046) | 0.093 | 0.000001<br>(0.079) | 0.5 |
| N | 3,1396 |  | 16,109 |  |

*Note*: NEB = number of children ever born, AFB = age at first birth, 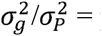 proportion of observed variance in the outcome associated with genetic variance across populations and sexes, 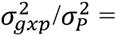 proportion of observed variance in the outcomes associated with *additional* genetic variance within populations, 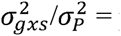 proportion of observed variance associated with *additional* genetic variance within sexes, 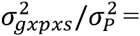 proportion of observed variance associated with *additional* genetic variance within populations and sexes, p-values are based on likelihood-ratio test comparing the full model with the model with one constraining the particular effect to be zero. All analyses include the first 20 Principal Components, and outcomes are standardized for sex, birth year and country. The model extends Model specification 2 (see Material & Methods) with **A**_**gxs**_,which is a matrix only with values for pairs of individuals within the same sex and **A**_**gxpxs**_,which is a matrix only with values for pairs of individuals with both the same sex and from the same population (following the same systematic as the extension of Model 3 to Model 4)

## Text SI1. additional Acknowledgment

The **Atherosclerosis Risk in Communities Study (ARIC)** is carried out as a collaborative study supported by National Heart, Lung, and Blood Institute contracts HHSN268201100005C, HHSN268201100006C, HHSN268201100007C, HSN268201100008C, HHSN268201100009C, HHSN268201100010C, HSN268201100011C, and HHSN268201100012C). The authors thank the staff and participants of the ARIC study for their important contributions. Funding for CARe genotyping was provided by NHLBI Contract N01-HC-65226. Funding for GENEVA was provided by National Human Genome Research Institute grant U01HG004402 (E. Boerwinkle).

**HRS** is supported by the National Institute on Aging (NIA U01AG009740). The genotyping was funded separately by the National Institute on Aging (RC2 AG036495, RC4 AG039029). Our genotyping was conducted by the NIH Center for Inherited Disease Research (CIDR) at Johns Hopkins University. Genotyping quality control and final preparation of the data were performed by the Genetics Coordinating Center at the University of Washington. Genotype data can be accessed via the database of Genotypes and Phenotypes (dbGaP, http://www.ncbi.nlm.nih.gov/gap, accession number phs000428.v1.p1). Researchers who wish to link genetic data with other HRS measures that are not in dbGaP, such as educational attainment, must apply for access from HRS. See the HRS website (http://hrsonline.isr.umich.edu/gwas) for details.

The **LifeLines** Cohort Study, and generation and management of GWAS genotype data for the LifeLines Cohort Study is supported by the Netherlands Organization of Scientific Research NWO (grant 175.010.2007.006), the Economic Structure Enhancing Fund (FES) of the Dutch government, the Ministry of Economic Affairs, the Ministry of Education, Culture and Science, the Ministry for Health, Welfare and Sports, the Northern Netherlands Collaboration of Provinces (SNN), the Province of Groningen, University Medical Center Groningen, the University of Groningen, Dutch Kidney Foundation and Dutch Diabetes Research Foundation. We thank Behrooz Z. Alizadeh, Annemieke Boesjes, Marcel Bruinenberg, Noortje Festen, Pim van der Harst, Ilja Nolte, Lude Franke, Mitra Valimohammadi for their help in creating the GWAS database, and Rob Bieringa, Joost Keers, René Oostergo, Rosalie Visser, Judith Vonk for their work related to data-collection and validation. The authors are grateful to the study participants, the staff from the LifeLines Cohort Study and the contributing research centers delivering data to LifeLines and the participating general practitioners and pharmacists. All data and samples collected by LifeLines are available to scientific researchers worldwide. It is also possible to prospectively collect additional data and samples in a selected group of LifeLines participants in an add-in study. Researchers can apply for data, samples or an add-on study by filling in the application form for research and submitting the completed form through our data catalogue, together with a selection of the requested data. Please contact dr. Salome Scholtens at, when you may need more specific information.

**QIMR** - Funding was provided by the Australian National Health and Medical Research Council (241944, 339462, 389927, 389875, 389891, 389892, 389938, 442915, 442981, 496739, 552485, 552498), the Australian Research Council (A7960034, A79906588, A79801419, DP0770096, DP0212016, DP0343921), the FP-5 GenomEUtwin Project (QLG2-CT-2002-01254), and the U.S. National Institutes of Health (NIH grants AA07535, AA10248, AA13320, AA13321, AA13326, AA14041, DA12854, MH66206). A portion of the genotyping on which the QIMR study was based (Illumina 370K scans) was carried out at the Center for Inherited Disease Research, Baltimore (CIDR), through an access award to the authors’ late colleague Dr. Richard Todd (Psychiatry, Washington University School of Medicine, St Louis). Imputation was carried out on the Genetic Cluster Computer, which is financially supported by the Netherlands Scientific Organization (NWO 480-05-003). N.W.H.M was supported by a PhD scholarship from the ANZ trust. S.E.M., is supported by the Australian Research Council (ARC) Fellowship Scheme. Dale R. Nyholt is supported by the Australian Research Council (ARC) Future Fellowship (FT0991022) and National Health and Medical Research Council (NHMRC) Research Fellowship (APP0613674) Schemes. The funders had no role in study design, data collection and analysis, decision to publish, or preparation of the manuscript. Researchers interested in using QIMR data can contact Nick Martin and Sarah Medland.

**STR (Swedish Twin Registry)** – The Jan Wallander and Tom Hedelius Foundation (P2012-0002:1), the Ragnar Söderberg Foundation (E9/11), The Swedish Research Council (421-2013-1061), the Ministry for Higher Education, The Swedish Research Council (M-2205-1112), GenomEUtwin (EU/QLRT-2001-01254; QLG2-CT-2002-01254), NIH DK U01-066134, The Swedish Foundation for Strategic Research (SSF). Researchers interested in using STR data must obtain approval from the Swedish Ethical Review Board and from the Steering Committee of the Swedish Twin Registry. Researchers using the data are required to follow the terms of an Assistance Agreement containing a number of clauses designed to ensure protection of privacy and compliance with relevant laws. For Further information, contact Patrik Magnusson.

The **TwinsUK** study was funded by the Wellcome Trust; European Community’s Seventh Framework Programme (FP7/2007–2013). The study also received support from the National Institute for Health Research (NIHR)- funded BioResource, Clinical Research Facility and Biomedical Research Centre based at Guy’s and St Thomas’ NHS Foundation Trust in partnership with King’s College London. SNP Genotyping was performed by The Wellcome Trust Sanger Institute and National Eye Institute via NIH/CIDR.

